# Molecule guided laser ablation as a novel therapeutic strategy to control itch

**DOI:** 10.1101/225391

**Authors:** Linda Nocchi, Mariangela D’Attilia, Nainika Roy, Rahul Dhandapani, Andrei Traista, Mariano Maffei, Laura Castaldi, Emerald Perlas, Paul A. Heppenstall

**Affiliations:** EMBL Rome, Via Ramarini 32, Monterotondo 00015, Italy; Molecular Medicine Partnership Unit (MMPU), Heidelberg, Germany

## Abstract

Itch is a major symptom of many chronic skin diseases that can exacerbate inflammation by provoking excessive scratching and causing skin damage. Here we develop a novel technology to control itch through molecular guided delivery of a phototoxic agent and near infrared (IR) illumination of the skin. Exploiting the selective binding of the pruritogen Interleukin-31 to itch sensing cells, we generate an engineered IL31^SNAP^ ligand derivative (IL31^K138A-SNAP^) that binds to cells but does not evoke signaling or provoke scratching when injected in vivo. Conjugation of IL31^K138A-SNAP^ to the photosensitizer IRDye®700DX phthalocyanine (IR700) and injection of the skin results in long-term reversal of scratching behavior evoked by IL31 upon near IR illumination. We further develop a topical preparation of IL31 ^K138A-SNAP^-IR700 that strikingly, reverses behavioral and dermatological indicators in mouse models of Atopic Dermatitis (AD) and the genetic skin disease Familial Primary Localized Cutaneous Amyloidosis (FPLCA). We demonstrate that this therapeutic effect results from selective retraction of itch sensing neurons in the skin, breaking the cycle of itch and disruption of the skin’s barrier function. Thus, molecule guided photoablation is a powerful new technology for controlling itch and treating inflammatory skin diseases.

## Introduction

Itch is a cutaneous sensory perception defined by the behavioral response it elicits: an urgent need to scratch ^1^. When itching becomes pathological, it can be irritating and distressful and have a dramatic impact on quality of life ^2^ Chronic itch generates a recurrent cycle whereby the more the skin is scratched, the more it itches ^3^ In turn, this may lead to serious damage to the skin barrier and thereby an increased risk of infection ^3^ There are many itch-associated diseases such as Atopic Dermatitis (AD) and psoriasis that respond poorly to current therapies ^4^ Identifying novel strategies to reduce itching is therefore critical and requires a deeper understanding of the underlying mechanisms.

Although several key molecules involved in transducing itch sensation have recently been described, the cellular and molecular mechanisms that drive chronic itch are not fully understood. The transduction of itch begins in the skin where a network of different cell types, such as keratinocytes, sensory nerves and immune cells respond to exogenous or endogenous pruritogens and initiate the cascade which ends with the scratch response ^5^ Two major itch pathways have been described: the histaminergic pathway which is mediated by histamine ^6^ and the non-histaminergic pathway which includes a plethora of other itch mediators, such as chloroquine, neuropeptides and inflammatory cytokines ^7^ It is becoming increasingly apparent that anti-histaminergic drugs are often ineffective against many chronic itch conditions such as Atopic Dermatitis and psoriasis ^8, 9^ Attention has now turned to non-histaminergic pathways to control itch sensation under pathological conditions ^10^

Amongst anti-histamine resistant itch mediators, the cytokine Interleukin 31 (IL-31) has recently attracted much attention as a novel target molecule for chronic itch therapy ^11^. IL-31 is a cytokine with a prominent skin tropism that is produced preferentially by T helper-type 2 cells ^12^ It signals through a heterodimeric receptor composed of IL31RA and OSMR which are expressed in epithelial cells, keratinocytes and sensory neurons ^13, 14^ Transgenic mice overexpressing IL-31 develop severe pruritus, alopecia and skin lesions that resemble lesioned skin from patients with atopic dermatitis ^11^. Moreover, numerous studies have reported an association of IL-31 with inflammatory skin diseases with a severe pruritic component such as atopic dermatitis, prurigo nodularis, allergic contact dermatitis, and cutaneous T-cell lymphoma ^15-18^ For example, IL-31 mRNA is up-regulated in human patients and mouse models of atopic dermatitis ^15, 19^ and increased levels of IL31 correlate with disease severity ^20, 21^. A common IL31 haplotype has also been associated with non-atopic eczema in three independent European populations, marking this as the first genetic risk factor for non-atopic eczema ^22^ Furthermore, mutations in the IL31 receptors IL31RA and OSMR underlie Familial Primary Localized Cutaneous Amyloidosis (FPLCA), a hereditary skin disorder characterized by the deposition of amyloid in the dermal papilla and severe pruritus ^23-27^ Intriguingly, it has been suggested that the major pathology evoked by IL-31 is to induce pruritus, rather than directly causing skin lesions per se ^28^ Thus, therapies which reduce scratching and break the cycle of itch and disruption of the skin’s barrier function ^29^ may be the most effective strategies for improving the quality of life for patients with chronic pruritic disease ^3^ Indeed, recent phase I and 2 clinical trials of a humanized antibody against IL31RA (Nemolizumab, CIM331) have shown some promise in improving pruritus in patients with moderate to severe atopic dermatitis ^30, 31^.

Here we have asked whether the IL31 ligand may serve as an alternative means of accessing pathways in the skin which provoke itch. Because IL31 receptors are expressed by itch sensing cells, we reasoned that conjugation of the IL31 ligand to a photosensitizer may allow for specific delivery of phototoxic complexes to these cells in the skin, and thus enable light activated suppression of itch sensation at its source. To achieve this aim we generated a recombinant IL31 fusion protein with a SNAP tag placed at its C-terminus that would allow for its chemical modification with benzylguanine (BG) derivatized probes. We demonstrate that IL31^SNAP^ is functional and can be conjugated to a photosensitizer to control IL31-induced scratching behavior. We further identify an IL31 mutant (IL31^K138A^) that binds to keratinocytes but no longer provokes signaling or scratching. Delivery of IL31^K138A-SNAP^- IR700 to the skin and subsequent near infrared illumination inhibits acute scratching and reverses behavioral and dermatological indicators in mouse models of Atopic Dermatitis and FPLCA. Intriguingly, this effect appears to be dependent upon retraction of sensory fibers from the epidermis of the skin, thus breaking the cycle of itch and disruption of the skin’s barrier function.

## Results

### Generation and characterization of IL31^SNAP^

Recombinant IL31 was produced with a C-terminal fusion of SNAP (IL31^SNAP^) in E. Coli. Following purification and refolding from inclusion bodies, IL31^SNAP^ was efficiently labelled with BG derivatized fluorophores (Figure 1a) indicating that the SNAP tag was successfully expressed and correctly folded in the fusion protein. To determine whether IL31^SNAP^ was functional we first performed binding studies in primary keratinocyte cultures. IL31^SNAP^ was labelled in vitro with BG-Surface549 and applied to keratinocytes from wildtype or IL31 Receptor A (IL31 RA) knockout mice (IL31 RA^-/-^). At a range of concentrations we observed strong fluorescent signal internalized in wildtype keratinocytes that was not present in cells for IL31RA^-/-^ mice (Figure 1b and c and Supplementary Figure 1a-k). We further examined whether IL31^SNAP^ was active by quantifying scratching behavior in mice upon intradermal injection. IL31^SNAP^ evoked robust scratching that was comparable in duration and intensity to native recombinant IL31 in wildtype mice (Figure 1d and Supplementary Figure 1l). In IL31RA^-/-^ mice, IL31^SNAP^ and IL31 did not evoke scratching (Figure 1d). Thus, the IL31^SNAP^ retains the functional properties of native IL31.

**Figure 1.**
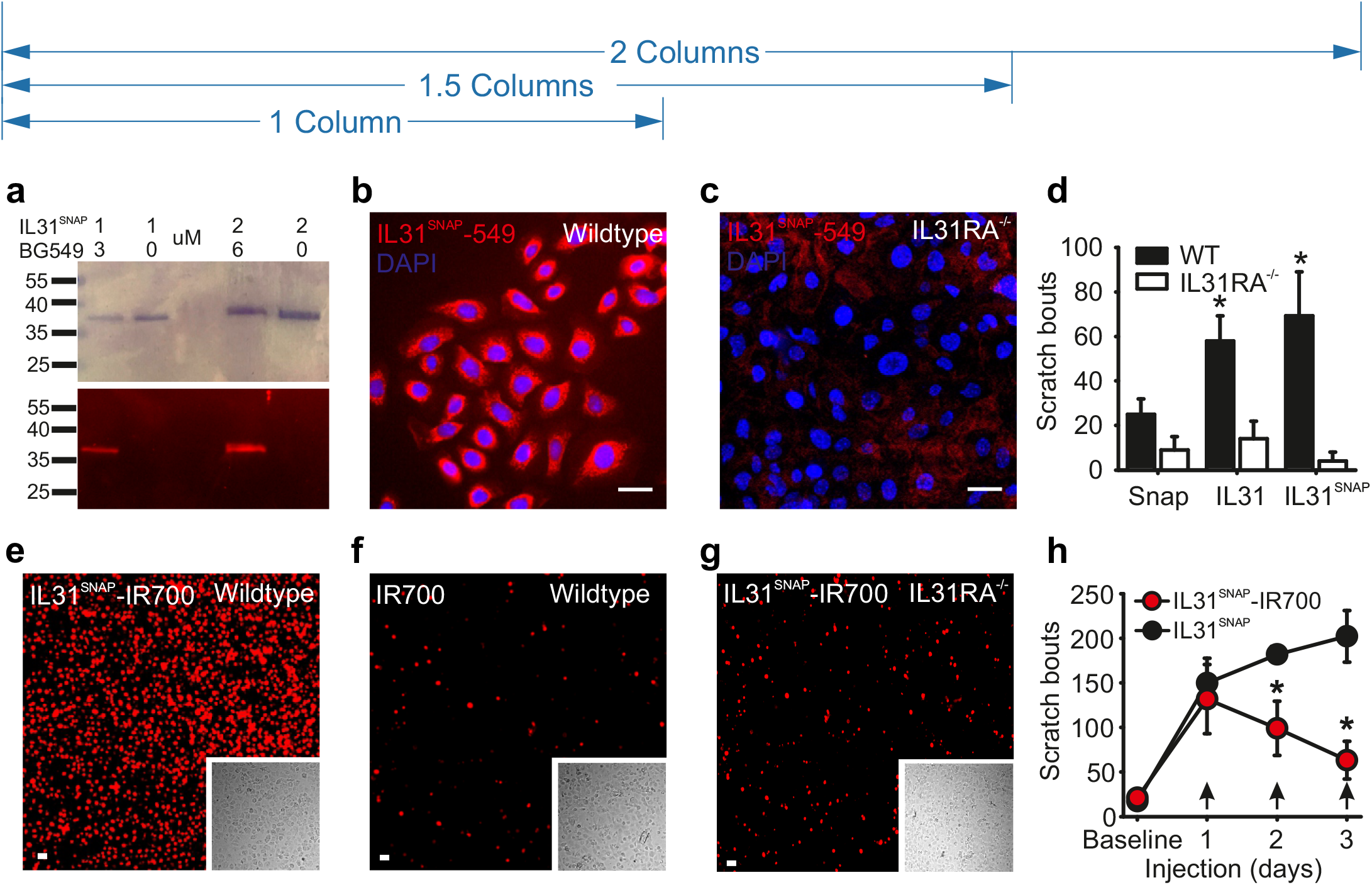
IL31^SNAP^ labelling and photoablation. (**a**) Representative Coomassie (upper panel) and fluorescence (lower panel) gel showing the binding of IL31^SNAP^ with the fluorescent substrate BG549 at a 1:3 molar ratio. First and fourth lanes represent the binding of 1 and 2 μM IL31^SNAP^ respectively, with BG549. Second and last lanes represent the protein IL31^SNAP^ alone, 1 and 2 μM respectively. (**b**) Primary keratinocyte culture from wildtype and (**c**) IL31RA^-/-^ mice labelled with IL31^SNAP^ + BG549 (in red). Nuclei were stained with Dapi (in blue). Scale bar 20 μm. (**d**) Scratching evoked by intradermal injection of SNAP, IL31 or IL31^SNAP^ in the nape of the neck of wildtype (WT, n=4, black bar) and IL31 RA^-/-^ mice (n=4, white bar). * p<0.05 (t-Test). (e, f, g) Propidium Iodide staining to assess cell death (in red) 24 hours after photoablation was performed on primary wildtype keratinocytes labelled with IL31^SNAP^-IR700 (**e**), with IR700 only (**f**), or IL31RA^-/-^ keratinocytes labelled with IL31SNAP-IR700 (**g**). Insets represent the corresponding brightfield images. Scale bar 50 μm. (**h**) Scratching behavior evoked by 3 days of injection (arrows) with IL31^SNAP^ (n=4, black circle) or IL31^SNAP^ + IR700 (n=4, red circle) with near IR illumination. Baseline refers to spontaneous scratching before the first injection. * p<0.05 (Bonferroni PostHoc test). Error bar indicate standard error (SEM).

### IL31 mediated photoablation

To manipulate IL31 receptor expressing cells in vivo, we reasoned that IL31^SNAP^ may allow for targeted photoablation of these cells through delivery of a photosensitizing agent. We synthesized a benzylguanine modified derivative of the highly potent near-infrared photosensitizer IRDye®700DX phthalocyanine (IR700) and conjugated it in vitro to IL31^SNAP^ 32 Application of IL31 ^SNAP^-IR700 to keratinocytes followed by 1 minute illumination provoked substantial cell death (Figure 1e) that was not evident in keratinocytes only treated with IR700 + illumination (Figure 1f) or in keratinocytes from IL31RA^-/-^ mice (Figure 1g). We further examined photosensitizer effects in vivo by injecting IL31 ^SNAP^-IR700 subcutaneously, applying near infrared light to the skin, and monitoring IL31 evoked scratching behavior. Strikingly, a progressive decrease in scratching bouts was observed when IL31 ^SNAP^-IR700 was injected for three consecutive days and the skin illuminated (Figure 1h).

### Generation and characterization of a non-signaling IL31 mutant

A conceptual problem of using IL31^SNAP^ therapeutically is that it in itself evokes itch. We therefore sought to engineer IL31^SNAP^ to obtain a ligand that still binds to IL31 receptor complex but no longer promotes signaling. From a previous structural study ^33^ we selected an IL31 point mutant IL31^K134A^ that was reported to exhibit reduced signaling in cells expressing human IL31 receptors. We generated a recombinant mouse IL31^K138A-SNAP^ fusion protein (orthologous to human IL31^K134A^), labelled it with BG-Surface549 and applied it to keratinocytes. Pronounced fluorescence was evident in cells from wildtype mice treated with fluorescent IL31 ^K1^3^8A-SNAP^ (Figure 2a), at a similar concentration range to that observed with IL31^SNAP^ (Supplementary Figure 2a-k). Importantly, such signal was not present in IL31RA^-/-^ keratinocytes (Figure 2b). We further assessed cellular signaling pathways activated by IL31^SNAP^ and IL31^K138A-SNAP^ in the skin by examining levels of phosphorylated Akt, pMAPK and pSTAT3 which have all been previously implicated in IL31 downstream signaling ^14, 33, 34^ Mice were injected subcutaneously with IL31^SNAP^ and IL31^K138A-SNAP^ and skin harvested 1 hour later for immunoblot analysis. We observed an increase in phosphorylation of the downstream pathways in skin injected with IL31^SNAP^ and this increase was absent in skin injected with IL31^K138A-SNAP^ (Figure 2c and d). As a final test for the functional activity of IL31^K138A-SNAP^, we assayed its capacity to provoke scratching behavior in mice. In contrast to IL31^SNAP^ which induced robust scratching when injected intradermal, IL31^K138A-SNAP^ did not evoke any scratching above baseline levels in mice (Figure 2e). Thus, the engineered ligand IL31^K138A-SNAP^ may offer a powerful means of targeting cells involved in itch without triggering itch in itself.

**Figure 2.**
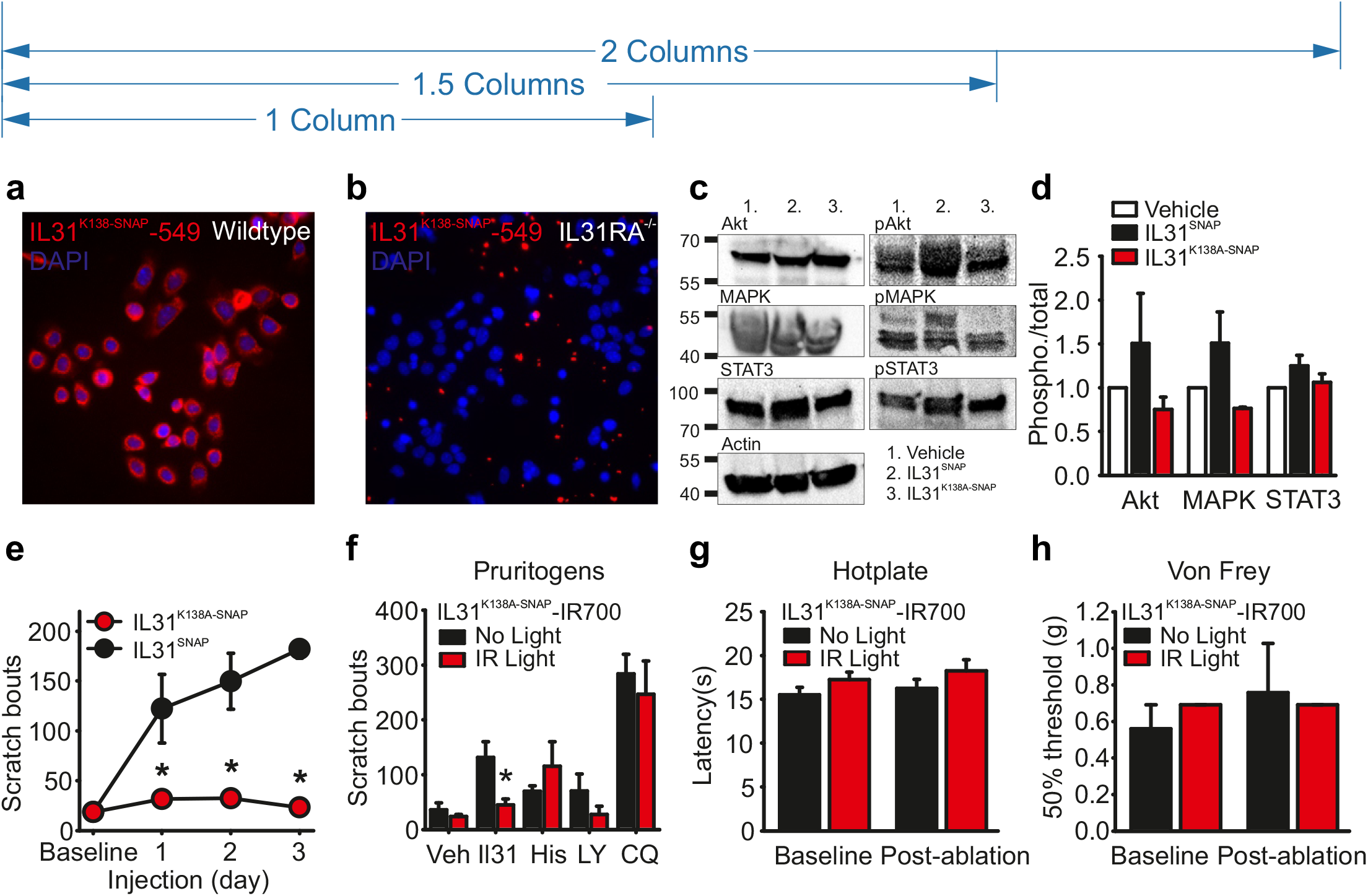
Functional analysis of IL31^K138A-SNAP^. (**a**) Primary keratinocyte cultures from wild type and (**b**) IL31 RA^-/-^ mice labelled with IL31^K138A-SNAP^ + BG549 (in red). Nuclei were stained with DAPI (in blue). Scale bar 20 μm. (**c**) Representative western blots showing the expression level of AKT, phospho AKT, MAPK, phospho-MAPK, STAT3, phospho STAT3 and Actin (loading control) in skin injected with vehicle (PBS, lane 1), IL31^SNAP^ (lane 2) and IL31^K138A-SNAP^ (lane 3). (**d**) Levels of each protein were expressed as the ratio between the phosphorylated form and the total counterpart and then normalized to the vehicle-treated sample. (**e**) Scratching response evoked by 3 consecutive days of injection of IL31^SNAP^ (n=4, black circle) and IL31^K138A-SNAP^ (n=8, red circle). * p<0.05 (Two-Way Anova followed by Bonferroni test). (**f**) Scratching response evoked by the injection of vehicle (PBS, n=4) and different pruritogens (IL31, Histamine, LY344864, and Chloroquine CQ) after mice were injected for 3 consecutive days with IL31^K138A-SNAP^-IR700, with (n=5, red bar) and without (n=4, black bar) near IR illumination. * p<0.05 (t-Test). (**g**) Hot Plate test after 3 days of injection of IL31^K138A-SNAP^-IR700 into the hind paw of the mice, with (n=4, red bar) and without (n=4, black bar) near IR illumination. Baseline refers to the thermal latency before the first injection was performed. Bar graphs represent the latency expressed in seconds (**s**) of the paw withdrawal in response to heat. (**h**) von Frey test after 3 days of injection of IL31^K138A-SNAP^-IR700 into the hind paw of the mice, with (n=4, red bar) and without (n=4, black bar) near IR illumination. Baseline refers to the mechanical threshold before the first injection was performed. Bar graphs represent the force expressed in grams (**g**) required to trigger a 50% response. Error bars indicate SEM. Baseline refers to spontaneous scratching before the first injection. Number of scratching bouts was counted over a 30 minutes recording time.

### IL31^K138A-SNAP^-IR700 mediated photoablation and acute itch

To characterize IL31^K138A-SNAP^ mediated photoablation in vivo, we first tested its efficacy at alleviating IL31 provoked itch. Mice were treated for three consecutive days with IL31^K138A-SNAP^-IR700 and the skin was illuminated with near-IR light. Strikingly, IL31-induced scratching behavior was abolished in these animals (Figure 2f), and this persisted throughout an 8 week observation period (Supplementary Figure 2l). In control mice that received an IL31^K138A-SNAP^-IR700 injection but were not illuminated, we observed no reduction in IL31-evoked scratching (Figure 2f). We next tested the effects of photoablation on scratching provoked by other acutely applied pruritogens. Intriguingly, IL31^K138A-SNAP^ guided photoablation had no significant effect on Histamine, LY344864 (a serotonin 5-HT1F receptor agonist) or Chloroquine evoked itch (Figure 2f). Finally, to assess the specificity of photoablation to itch sensation, we examined other sensory modalities after treatment. Using the Hot Plate test to assay thermal sensitivity (Figure 2g) and calibrated von Frey filaments to measure mechanical sensitivity (Figure 2h), we observed no difference in response properties of treated mice, indicating that IL31 guided laser ablation is indeed a selective and effective means of disrupting IL31 evoked itch.

### IL31^K138A-SNAP^-IR700 mediated photoablation and Atopic Dermatitis

We examined the effects of IL31^K138A-SNAP^-IR700 ablation on inflammatory itch using the Calcipotriol mouse model of Atopic Dermatitis ^35^ To assess the effectiveness of treatment we monitored three indicators of clinical progression: scratching behavior, skin integrity and skin histology. We first determined whether pretreatment with IL31^K138A-SNAP^-IR700 would abolish the development of the disease, and then investigated whether post-treatment, upon establishment of inflammation, could reverse symptoms.

As previously reported ^35^, application of Calcipotriol to skin provoked a severe Atopic Dermatitis-like phenotype that was evident as a progressive increase in the number of spontaneous scratching bouts over time, distinct skin damage, and a thickening of the epidermis and cell infiltration (Supplementary Figure 3a-d). Injection of IL31^K138A-SNAP^-IR700 and subsequent near IR illumination of the skin for 3 days prior to Calcipotriol application completely abolished the development of all indicators in this model. Thus, scratching behavior remained at baseline levels (Figure 3a), skin thickness and histological characteristics were not altered (Figure 3b and c), skin appeared healthy and typical features of dermatitis such as redness and scaling were entirely absent (Figure 3d). To control for a pharmacological antagonistic effect of IL31^K138A-SNAP^ we performed identical experiments in the absence of near IR illumination and observed the normal development of dermatitis-like symptoms (Figures 3a-d). Similarly, near IR light and IR700 alone were also ineffective at blocking the progression of the condition (Supplementary Figure 3e-h).

**Figure 3.**
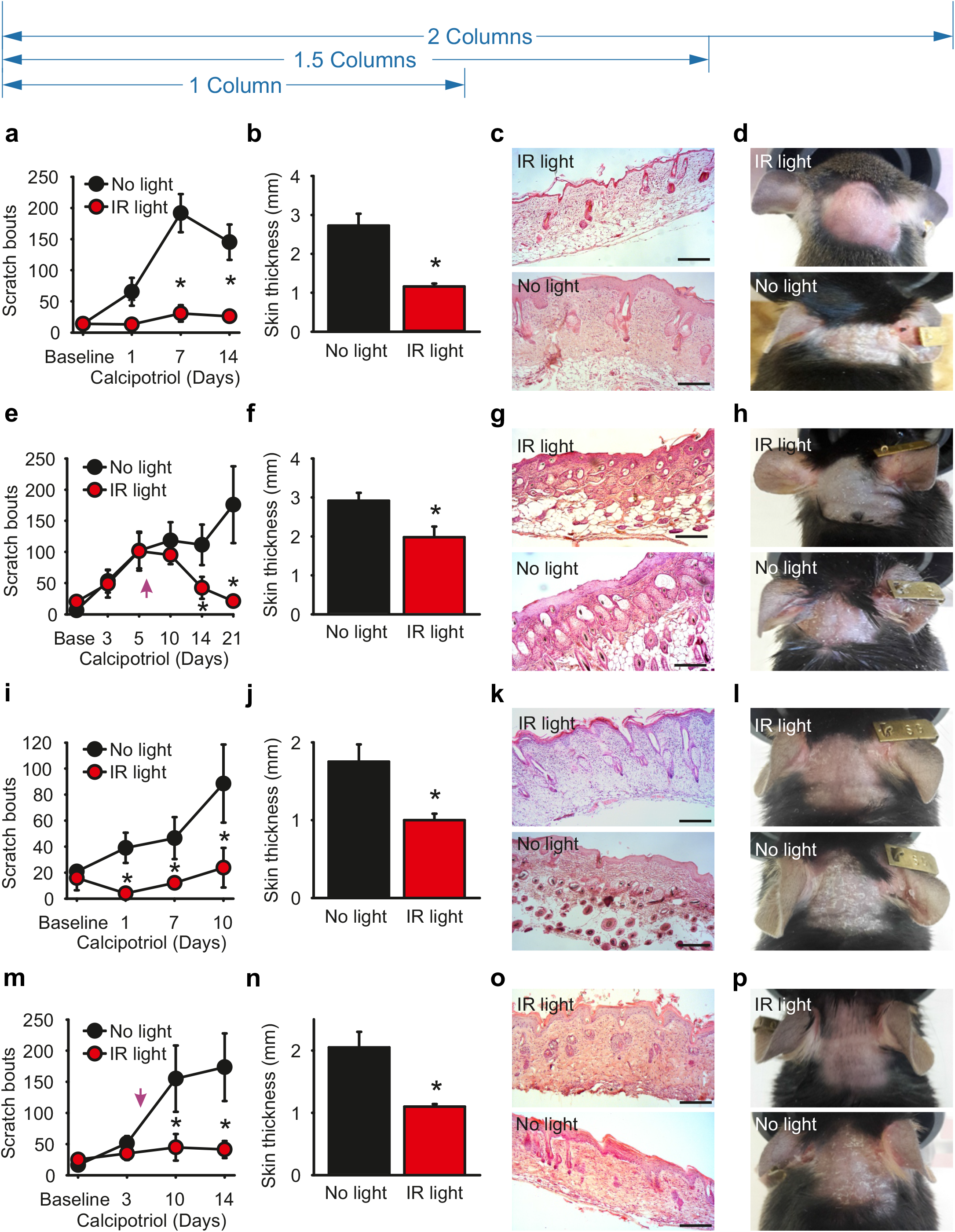
IL31^K138A-SNAP^ guided photoablation prevents and reverses atopic dermatitis-like symptoms. **a-d**). Prevention of atopic dermatitis-like symptoms (**a**) Number of scratch bouts in response to 14 days of Calcipotriol treatment in mice pre-injected with IL31^K138A-SNAP^-|R700, with (red circle, n=7) or without (black circle, n=8) near IR illumination. * p<0.05 (Two-Way Anova followed by Bonferroni test). (**b**) Skin thickness was measured at day 14 of Calcipotriol treatment * p<0.05 (t-Test). (**c**) Representative Hematoxylin and Eosin staining of AD skin sections of mice treated with (Top panel) and without (Lower panel) near IR illumination. (**d**) Representative skin pictures of AD mice with (Top panel) and without (Lower panel) near IR treatment. (e-h) Rescue of atopic dermatitis-like symptoms (**e**) Scratching bouts in response to 21 days of Calcipotriol treatment of mice treated (n=6, red circle), or not (n=6, black circle) with near IR illumination. Arrow indicates the day of the first photoablation. * p<0.05 (Two-Way Anova followed by Bonferroni test). (**f**) Skin thickness at day 21 of Calcipotriol treatment. * p<0.05 (t-Test). (**g**) Representative Hematoxylin & Eosin staining of skin sections of AD mice with (Top panel) and without (Lower panel) near IR illumination. (**h**) Representative skin pictures of AD mice with (Top panel) and without (Lower panel) near IR treatment (i-l) Prevention of atopic dermatitis-like symptoms using a topical delivery system. (**i**) Scratching bouts in response to 10 days of Calcipotriol in mice pretreated topically with IL31^K138A-SNAP^-IR700, with (n=4, red circle) and without (n=4, black circle) near IR illumination. * p<0.05 (Two-Way Anova followed by Bonferroni test). (**j**) Skin thickness after 14 days of Calcipotriol treatment. * p<0.05 (t-Test). (**k**) Representative Hematoxylin & Eosin staining of skin sections after 10 days of Calcipotriol treatment. (**l**) Representative skin pictures. (m-p) Rescue of atopic dermatitis symptoms using topical delivery (**m**) Scratching behavior evoked by 14 days of Calcipotriol in mice topically treated with IL31^K138A-SNAP^-IR700, with (n=7, red circle) and without (n=8, black circle) near IR illumination. Arrow indicates the first day of photoablation treatment. (**n**) Skin thickness at day 14 of Calcipotriol treatment. * p<0.05 (t-Test). (**o**) Representative Hematoxylin & Eosin staining of skin sections after 14 days of Calcipotriol (**p**) Representative skin pictures. Error bars indicate SEM, scale bars 200μm.

For IL31^K138A^-guided photoablation to have therapeutic relevance, it must also be effective in reversing already established skin inflammation. We therefore treated mice with Calcipotriol for 7 days until severe symptoms were evident. IL31^K138A-SNAP^-IR700 was then injected subcutaneously and near IR light applied to the skin for three consecutive days. Strikingly, we observed a rescue of all disease indicators over the course of 1 week. Thus, scratching behavior returned to baseline levels (Figure 3e), and skin thickness (Figure 3f), morphology (Figure 3g) and structure (Figure 3h) became indistinguishable from healthy mice. Such reversal of dermatitis-like symptoms was not evident in control experiments where IL31^K138A-SNAP^ was applied without subsequent near IR illumination (Figure 3e-h).

To further improve the clinical applicability of IL31^K138A^ guided photoablation, we sought to develop a formulation that would allow for topical, pain-free application of IL31^K138A-SNAP^- IR700. We selected a water-in-oil micro-emulsion preparation based upon previous evidence that this type of formulation can deliver high molecular weight proteins across the dermal barrier ^36^ IL31^K138A-SNAP^-IR700 was loaded into the aqueous phase of the microemulsion, applied topically, and 20 minutes later skin was illuminated with near IR light. Similar to subcutaneous delivery, topical application of IL31^K138A-SNAP^-IR700 both prevented (Figure 3i-l) and reversed (Figure 3m-p) Calcipotriol provoked dermatitis-like symptoms. This was evident as a return to baseline levels of scratching behavior (Figures 3i and m) and a normalization of skin structure and histology (Figures 3j-l and n-p). Thus, molecule guided delivery of a photosensitizer complex allows for on-demand, pain-free control of inflammatory itch.

### Generation of a mouse model of the genetic skin disorder Familial Primary Localized Cutaneous Amyloidosis (FPLCA)

FPLCA is a rare, hereditary skin disorder driven by mutations in the IL31 receptors IL31RA and OSMR, and characterized by severe pruritus ^37^ We sought to generate a mouse model of FPLCA to further test the therapeutic potential of IL31^K138A-SNAP^ photoablation in a more chronic and challenging clinical setting. We used CRISPR/Cas9 technology to knock-in an S476F mutation into the Il31ra locus, orthologous to the human FPLCA mutation IL31 RA^S521F 26^ From 14 offspring, we obtained one IL31 RA^-/-^ mouse and one IL31 RA^S476F^ mouse that were confirmed by sequencing (Supplementary Figure 4b). Upon expansion of the IL31RA^S476F^ line we observed that both heterozygote and homozygous mice spontaneously developed phenotypes that resembled symptoms of FPLCA. These included a significant increase in spontaneous scratching behavior (Supplementary Figure 5a), deposition of Congo red positive amyloid in the skin (Supplementary Figure 5c and d), thickening of the epidermis, and cellular infiltration of the dermis (Supplementary Figures 5e-h). Of note, we also observed that IL31^SNAP^ (Supplementary Figure 5b) and IL31^K138A-SNAP^ (Figure 4a) robustly labelled keratinocytes from IL31RA^S476F^ mice at similar concentrations to those used on wildtype cells.

**Figure 4.**
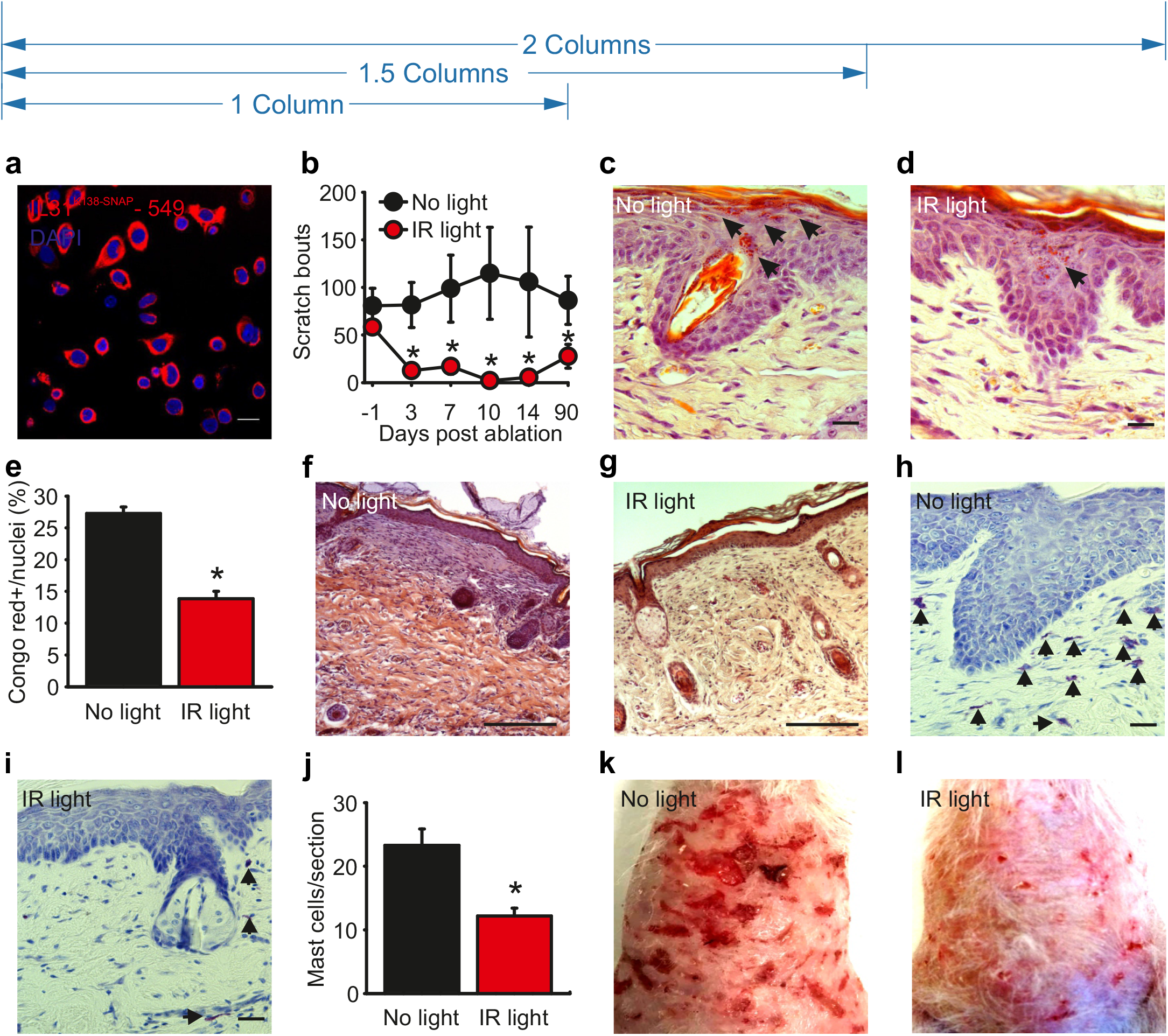
IL31^K138A-SNAP^ guided photoablation reverses indicators of FPLCA. (**a**) IL31^K138A-SNAP^+BG549 (in red) labels primary keratinocytes from IL31RA^S476F^ mice. Nuclei are stained with Dapi (in blue). Scale bar 20 μm. (**b**) Spontaneous scratching behavior in IL31 RA^S476F^ mice before any treatment (-1) and following topical application of IL31^K138A-SNAP^ – IR700, with (n=5, red circle) and without (n=4, black circle) near IR illumination. * p<0.05 (2-Way Anova followed by Tukey test). (c-d) Representative Congo Red staining for amyloid deposits in skin sections of IL31RA^S476F^ mice (n=3) treated topically with IL31^K138A-SNAP^ – IR700 without (**c**) or with near IR illumination (**d**); arrows indicate Congo red positive amyloid deposits. Scale bar 50 μm. (**e**) Quantification of Congo red staining expressed as percentage of the total number of epidermal cells. * p<0.001 (t-Test). (f-g) Representative Hematoxylin and Eosin staining of skin sections of IL31 RA^S476F^ mice (n=3) topically treated with IL31^K138A-SNAP^-IR700 without (**f**) or with illumination (**g**), showing reduced dermal infiltration of eosinophilic material upon photoablation treatment. Scale bar 200 μm. (h-i) Representative Toluidine blue staining for mast cells in skin sections of IL31RA^S476F^ mice treated topically with IL31^K138A-SNAP^ – IR700, without (**h**) and with (**i**) near IR illumination. Arrows indicate toluidine blue positive mast cells. Scale bar 50 μm. (**j**) Quantification of Toluidine blue staining expressed as number of positive mast cells per section (10 sections, n=2); * p = 0.002. (**k-l**) Representative images of skin of IL31 RA^S476F^ mice 90 days after they have been treated topically with IL31^K138A-SNAP^-IR700 in the absence of illumination (**k**) or with near IR illumination (**l**). All error bars indicate standard error (SEM).

**Figure 5.**
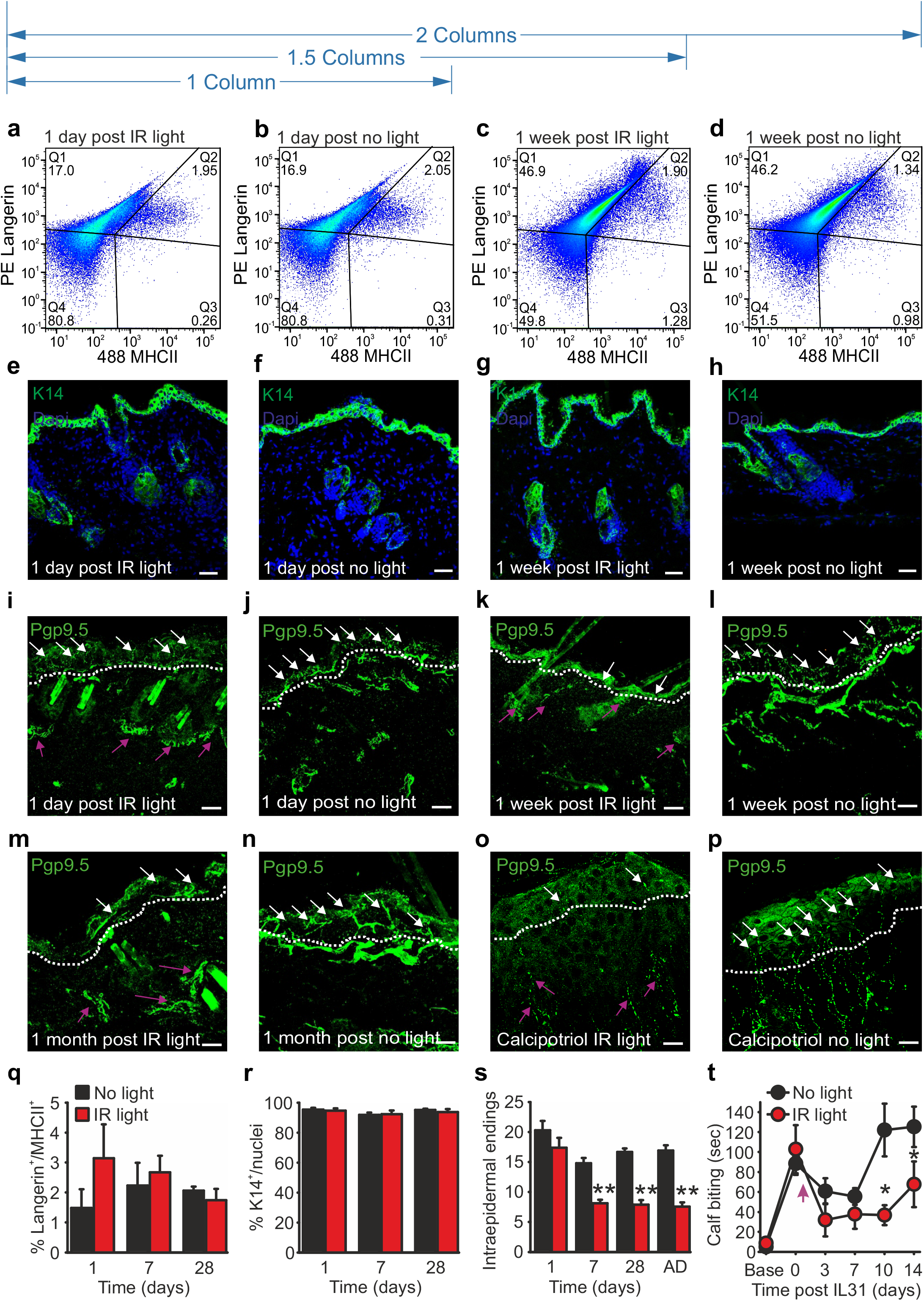
Epidermal free nerve endings are the cellular target of IL31^K138A-SNAP^ guided photoablation. (**a-d**) No loss of epidermal migratory dendritic cells upon IL31^K138A-SNAP^ mediated photoablation. Representative flow cytometry plots showing the percentage of cutaneous dendritic cells isolated from the back skin of mice at 1 day (**a, b**) and 1 week (**c, d**) after topical application of IL31^K138A-SNAP^-IR700, with (**a, c**) and without (**b, d**) near IR illumination. Q2 represents epidermal migratory dendritic cells (Langerhans cells, Langerin^+^/MHCII^+^), Q1 represents Langerin+/MHCII-resident inactivated dendritic cells, Q3 represents Langerin^-^/MHCII^+^ dermal migratory dendritic cells. Percentage of total events (10^5^ cells) are indicated in each box. (**e-h**) No loss of basal keratinocytes upon IL31^K138A-SNAP^ mediated photoablation. Representative immunofluorescence staining of back skin with a K14 antibody at 1 day (e,f) and 1 week (g,h) after topical application of IL31^K138A-^SNAP-IR700, with (**e, g**) and without (**f, h**) near IR illumination. (**i-p**) Loss of epidermal free nerve endings upon IL31^K138A-SNAP^ mediated photoablation. Representative immunofluorescence staining of Pgp9.5 positive nerve fibers innervating back skin at 1 day (**i,j**), 1 week (**k,l**), 1 month (**m,n**), or upon establishment of Atopic Dermatitis (**o,p**), after IL31^K138A-SNAP^-IR700 application. Dotted line represents the dermal-epidermal junction, white arrows indicate epidermal free nerve endings which are lost upon illumination, magenta arrows show innervations around hair follicles which are spared. (**q-s**) Quantification of the number of Langerhans cells (**q**), keratinocytes (**r**), or epidermal free nerve endings (**s**) at the indicated time point post IL31^K138A-SNAP^-IR700 application (** p<0.001, t-Test). (**t**) Peri-sciatic nerve application of IL31^K138A-SNAP^-IR700 reduces IL31 biting behavior upon near infrared illumination (n=5, red circle) compared with no light treated group (n=4, black circle). Arrow indicates the day of nerve photoablation. * p<0.05 (2-Way ANOVA followed byTukey test). Scale bars indicate 50 μm, error bars indicate SEM.

### IL31^K138A-SNAP^-IR700 mediated photoablation reverses symptoms of FPLCA

To assess whether targeted photoablation can reverse FPLCA symptoms we applied IL31^K1 38a-snap^-IR700 micro-emulsion topically to back skin of IL31RA^S476F^ mice. Strikingly, we observed a significant decrease in spontaneous scratching behavior upon illumination that became evident at 3 days after ablation and continued for the 90 day observation period (Figure 4b). We further characterized histological parameters of FPLCA at 14 days post treatment. All pathological indicators were reversed by photoablation including density of amyloid deposition (Figure 4c-e), infiltration of dermal eosinophilic material (Figure 4f and g), and mast cell counts (Figure 4h-j). Moreover, redness, scaling and superficial scaring of skin was visibly improved 14 days after treatment and continued for the 90 day observation period (Figure 4k and l). Importantly, in the absence of illumination, we observed no difference in behavioral or histological indicators of FPLCA, suggesting that the cream on its own does not induce any effect.

### IL31^K138A-SNAP^-IR700 mediated photoablation in a mouse model of psoriasis

To investigate further the scope of IL31^K138A^ mediated photoablation, we tested its effectiveness in an additional skin disorder, psoriasis, which is not believed to have a strong dependence upon IL31 signaling ^38^ We used the Aldara mouse model of psoriasis where Aldara cream, containing the Toll-like receptor 7-ligand Imiquimod, is applied to the skin. In this model, neither pretreatment nor posttreatment with IL31^K138A-SNAP^-IR700 and illumination altered scratching behavior provoked by Aldara cream (Supplementary Figure 6). Similarly, skin thickness, erythema, inflammation and integrity remained unaltered by photoablation as shown by the Psoriasis Area and Severity Index (PASI) (Supplementary Figure 6). Thus, IL31^K138A^ guided photoablation is likely ineffective in treating psoriasis.

### Mechanism of action of IL31^K138A-SNAP^- mediated photoablation

To investigate the mechanism underlying IL31^K138A-SNAP^-IR700 mediated photoablation we focused our attention on the three major cell populations which express IL31 receptor complexes in the skin; cutaneous dendritic cells (specifically Langerhans cells), keratinocytes and free nerve endings innervating the skin ^13, 39-41^. Again, we performed photoablation by applying IL31^K138A-SNAP^-IR700 to the back skin and then analyzed samples from illuminated or control non-illuminated skin at 1 day, 1 week or 1 month post treatment.

We first considered cutaneous dendritic cells. Skin cell suspensions were labelled with antibodies against Langerin and MHCII, and quantified by flow cytometry. We observed no difference in any of the populations of Langerin and MHCII positive cells (Langerin+/MHCII^-^ resident inactivated dendritic cells, Langerin+/MHCII+ Langerhans cells, Langerin^-^/MHCII+ dermal migratory dendritic cells) at any time point after ablation (Figures 5a-d, q, and Supplementary Figure 7a and b). Turning to keratinocytes, we labelled skin sections with an antibody against Keratin 14 and quantified cellular density. Again, we observed no difference in keratinocyte numbers at any point after ablation (Figures 5e-h, r, and Supplementary Figure 7c and d). In agreement with these measures we also detected no difference in the number of TUNEL positive cells at 1 day, 1 week and 1 month post treatment, indicating that illumination does not provoke additional apoptosis in skin (Supplementary Figure 7e-j).

We next examined innervation of the epidermis using the pan-neuronal antibody Pgp9.5 to label nerve endings in skin sections. At one day after treatment we observed no difference in the number of free nerve endings crossing the dermal epidermal junction (Figure 5i, j and s). However, at one week and one month post ablation, a marked reduction in epidermal innervation density was evident (Figure 5k-n and s). Similarly, photoablation also provoked a prominent reduction in numbers of epidermal nerve fibers in skin taken from Calcipotriol treated mice (Figure 5o-p and s). Importantly, the density of presumed non-pruriceptive fibers innervating hair follicles and dermal structures was not reduced by IL31^K138A-SNAP^- IR700 treatment, supporting behavioral observations that the approach is specific to itch. Finally, to determine whether nerve ablation alone (without influencing other cells in the skin) is sufficient to reduce scratching behavior, we performed photoablation directly on the proximal sciatic nerve. IL31^K138A-SNAP^-IR700 was applied to the sciatic nerve at mid-thigh level and scratching/biting behavior evoked by IL31 injection of the innervation territory in the calf was monitored. Upon illumination of the sciatic nerve we observed a significant reduction in IL31 evoked behavior that was not evident in non-illuminated control animals. Thus, IL31^K138A-SNAP^ guided photoablation triggers retraction of pruriceptive sensory neurons in the skin, thereby alleviating scratching and the chain of events that leads to inflammation.

## Discussion

Chronic itch is a common and often overlooked condition that is difficult to treat and exerts a substantial societal burden ^42, 43^ Here we describe a novel approach to control itch that is highly effective in mouse models of Atopic Dermatitis and the genetic skin disease FPLCA. Exploiting the natural selectivity of the IL31 ligand to its receptors on itch sensing cells in the skin, we design an IL31 derivative that enables delivery of a photosensitizer to itch sensors in vivo without causing itch in itself. Through illumination of treated skin, we are able to reverse behavioral and histological indicators of atopic dermatitis and FPLCA from a single treatment regime. We demonstrate that this is dependent upon photosensitizer induced retraction of itch sensing neurons from the epidermis, and thus silencing of the neuronal driver of scratching at its source in the skin.

Current therapeutic options for alleviating chronic itch rely on treatments that aim at controlling skin barrier function and inflammation rather than targeting itch itself ^44^ For example, in atopic dermatitis patients, initial treatment involves the use of topical moisturizers, emollients and corticosteroids, followed by systemic treatments such as oral immunosuppressants or UV therapy if disease worsens ^4, 43^ All of these approaches suffer from severe adverse effects, and a significant problem in treating chronic itch is continued adherence to therapy, with most patients abandoning treatments within a year ^44^ Recently, the first mechanism based targeted therapy for atopic dermatitis was approved in the US, a human monoclonal antibody against the interleukin-4 receptor alpha subunit named Dupixent ^45-47^ In addition, further clinical trials are also underway for a humanized antibody against IL31RA (Nemolizumab) ^30, 31^, highlighting the potential of biologics in treating itch. It is currently not known what the long term effectiveness and safety of these treatments will be, and the cost of repeated treatment is likely to restrict their use to only the more severe cases of atopic dermatitis ^45, 48^.

The photoablation approach we describe here has several conceptual differences to the aforementioned strategies for treating pruritic diseases. Firstly, photoablation targets cells rather than single molecules, thus bypassing the enormous molecular complexity that underlies itch. A major difficulty in selecting targets for new antipruritic medications is the molecular redundancy that is inherent to itch sensation. By inactivating the integrating unit of itch, the sensory neuron, rather than individual molecules in the itch pathway, we circumvent this redundancy. Secondly, our data indicates that a single treatment regime with photosensitizer and light allows for long-lasting alleviation of scratching, thus facilitating skin healing. Indeed we observed reversal of IL31-evoked itch that persisted throughout an 8 week observation period, and in the FPLCA model, behavioral and histological indicators were normalized for at least 3 months. In the Calcipotriol model of dermatitis we also observed recovery that continued for the maximal ethically acceptable observation period. Thirdly, through local, on-demand application of ligand-photosensitizer complex and light, we are able to achieve precise control of drug action, thus avoiding systemic side effects and allowing therapy to be tuned to disease severity. Further development could also allow for systemic delivery of the photosensitizer complex combined with local application of light. Finally, we have developed an engineered ligand, rather than an antibody or small molecule to target cells. We chose this approach because we wished to exploit the naturally high affinity of IL31 to its receptor, and because of ease of production of recombinant ligand. A similar approach could therefore by readily adapted to other cell surface receptors such as for example IL17 receptors in psoriasis, without the need for extensive screening to identify novel binders. Moreover, existing antibodies or small molecules that target membrane receptors could also be co-opted to deliver photosensitizers, analogous to delivery methods developed for photodynamic therapy of cancer ^49^

Beyond atopic dermatitis, IL31 has been implicated in a number of other pruritic diseases such as prurigo nodularis, cutaneous T-cell lymphoma, and chronic idiopathic pruritus ^15, 17, 18, 50^, and targeted photoablation could conceivably also have therapeutic efficacy in these disorders. To further examine the scope of IL31 mediated photoablation, we chose to focus on the rare genetic skin disease FPLCA, which is driven by mutations in the IL31 receptor complex ^24-27^ FPLCA is characterized by severe pruritus and localized deposition of amyloid in the dermal papilla ^23^, both of which are highly distressing and impact profoundly on the quality of life of patients. There are no effective treatment options for FPLCA and management of the disorder relies upon topical application of corticosteroids, UV phototherapy and systemic treatment with immunosuppressants ^23^. Here we describe the first mouse model of FPLCA, and demonstrate that mutant IL31RA^S476F^ mice develop phenotypes that are remarkably similar to human symptoms. Thus we observed a significant increase in spontaneous scratching behavior, damage and inflammation of the skin, and deposition of amyloid. We did not explore in detail the molecular mechanisms that underlie FPLCA, instead focusing on gauging the therapeutic potential of IL31^K138A^ mediated photoablation. Indeed we found that a single treatment regime led to a substantial improvement in disease progression that persisted throughout the 3 month observation period. We now expect that the IL31 RA^S476F^ mouse model will allow for further mechanistic investigations, and will also enable preclinical testing of other emerging IL31 orientated medications such as humanized IL31 RA antibodies ^30, 31^ and JAK inhibitors ^42, 51^ for treating FPLCA.

IL31 receptor complexes are expressed by cutaneous dendritic cells, keratinocytes, and free nerve endings in the skin ^13, 39-41^, and hypothetically, IL31^K138A^ mediated photoablation could induce cell toxicity in any of these cell types. Intriguingly, our analysis of these populations post ablation points towards a selective and sustained loss of neuronal innervation by IL31 ^K138A^-IR700. This raises the question as to why dendritic cells and keratinocytes are spared by the treatment. One possibility is that there is a transient loss of these cells which we are not able to detect because of ongoing cell turnover and re-infiltration of the skin. However, we did not observe any increase in TUNEL positive cells in the skin after photoablation suggesting that there is not a reduction in these cells through apoptosis. An alternative explanation is that sensory neurons, as excitable cells, are more susceptible to photosensitizer induced damage than dendritic cells or keratinocytes. Indeed, we have previously observed that exposure of sensory neurons to the photosensitizer fluorescein leads to a rapid loss of action potential firing and an increase in resting membrane potential ^52^, which could act as an initial trigger to promote neuronal toxicity.

Our data also place sensory neurons as a critical integration point in the transduction of inflammatory itch, and support growing evidence that neurons represent an attractive target for treating pruritic skin diseases ^1, 42, 53^ Of note, only a small proportion of sensory neurons express IL31 receptors ^54-56^, and together with previous genetic ablation experiments ^54^ our analysis indicates that these neurons are not overtly sensitive to acute application of other pruritogens such as histamine and chloroquine. Intriguingly, recent evidence indicates that these neurons are sensitized by activation of interleukin-4 receptor alpha (the target for the human monoclonal antibody Dupixent), and JAK1 signaling, and that this may promote their involvement in inflammatory itch ^42^ Indeed, atopic dermatitis is known to have a prominent neuronal component ^5, 57^, and it has also been argued the FPLCA could represent a neurodermatitis rather than a primary disorder of keratinoctyes^25^. Data presented here suggest that IL31 RA expressing sensory neurons are the key drivers of these disorders, and that their removal through photoablation is a powerful means of reversing disease progression.

Long-term elimination of the sensory nerve endings that signal itch could be considered a relatively excessive measure for treating pruritus. However, several other therapies share similar functional principles to IL31^K138A^ mediated photoablation but without the selectivity that ligand targeting offers. These include the use of psoralens as photosensitizers in UV therapy (PUVA) ^58^, and the topical application of capsaicin ^59-61^, both of which promote the retraction of nerve fibers in the skin ^62, 63^ These treatments are however associated with adverse effects such as burning pain, increased itchiness and erythema, limiting patient compliance ^58, 64^ Moreover, non-selective denervation may also lead to a loss of other sensations in addition to itch. By targeting directly a subset of pruriceptors using an engineered ligand and near infrared illumination we circumvent these issues, allowing for local on demand treatment of itch. Further development of the technology to target other membrane receptors and neuronal subsets may allow for treatment of a broad range of sensory disorders spanning itch and pain.

## Acknowledgments

We thank Pedro Moreira of EMBL Transgenic Services, Cora Chadick of EMBL Flow Cytometry Facility, Violetta Paribeni and Matteo Gaetani for technical support of our work. We also acknowledge the assistance of the Protein Expression and Purification Core Facility for the generation of IL31^SNAP^, IL31^K138A-SNAP^ and Cas9. This work was funded by EMBL and Fondazione Telethon.

## Methods

### Animals

Wild type, IL31RA^-/-^, and IL31RA^S476F^ mice were used at age of 8-10 weeks for all the behavioral studies. 1-3 day-old mice were used for primary keratinocyte culture. All mice were bred and maintained at the EMBL Neurobiology and Epigenetic Unit, Rome, in accordance with Italian legislation (Art. 9, 27. Jan 1992, no 116) under license from the Italian Ministry of Health, and in compliance with the ARRIVE guidelines. IL31RA^-/-^, and IL31 RA^S476F^ mice have been deposited in the European Mouse Mutant Archive.

### Production of recombinant IL31^SNAP^ and IL31^K138A-SNAP^

cDNAs for murine IL31 and SNAP tag were cloned into pETM-11 vector and expressed in E.Coli as fusion protein, carrying a His6-tag at the N-terminus. To generate the mutant IL31^K138A-SNAP^, site directed mutagenesis was performed according to the manufacturer’s instruction (Agilent, #200555). The proteins were isolated from inclusion bodies, solubilized, refolded, and purified using a Ni-NTA resin (Qiagen, #30210). Eluted fractions were then pooled, concentrated and stored for further analysis.

### Synthesis of BG-IR700

3mg of IRDye®700DX N-hydroxysuccinimide ester fluorophore (LI-COR Biosciences GmbH, Bad Homburg, Germany) were dissolved in 150μl DMSO and treated with 1.5mg BG-PEG11-NH2 and 5μl diisopropylethylamine. After 1h, the product BG-PEG11-IRDye®700DX was purified by HPLC using a Waters Sunfire Prep C18 OBD 5μM; 19 x 150 mm column using 0.1M triethylammonium acetate (TEAA)] (pH 7.0) and 0.1M TEAA in water/acetonitrile 3:7 (pH 7.0) as mobile phases A and B, respectively. A linear gradient from 100% A to 100% B within 30 minutes was used. The fractions containing the product were lyophilized.

### Primary Keratinocyte culture

Primary keratinocytes were isolated from 1-3 day-old wild type, IL31RA^-/-^ and IL31RA^S476F^ mice as previously described ^65^ Briefly, newborn mouse skin was removed and incubated flat in cold and freshly thawed trypsin overnight, at 4°C, with the dermis side down. The next day, epidermis was peeled off, triturated and keratinocytes were cultured in serum free media (Invitrogen #10744-019) on Collagen I (Sigma # 122-20) – coated dishes (Ibidi #81151). All experiments were performed on 48-72 hours cultured cells.

### *In-vitro* labeling

For keratinocyte labelling, 1 μM IL31^SNAP^ or IL31^K138A-SNAP^ was coupled with 3 μM BG549surf (NEB # S9112) for 1 hour at 37°C in CIB buffer (NaCl 140 mM; KCl 4 mM; CaCl2 2 mM; MgCl2 1 mM; NaOH 4.55 mM; Glucose 5 mM; HEPES 10 mM; pH 7.4). Cells from wild type or IL31 RA^-/-^ or IL31 RA^S476F^ mice were incubated with the coupling reaction for 15-20 minutes at 37°C, then washed 3 times in CIB. 1μg/ml Hoechst was used for nuclear staining. All images were visualized with a Leica SP5 confocal microscope.

### *In-vitro* photo-ablation

1μM IL31^SNAP^ and 3μM IR700 were coupled for 1 hour at 37°C. The coupling reaction was applied for 15-20 minutes at 37¼C on primary wild type and IL31RA^-/-^ keratinocytes. Cells were then exposed to near infra-red light (680nm) at 40J/cm^2^ for 1 minutes. 24 hours after light exposure cell death was assessed by Propidium Iodide (PI) staining (Invitrogen # P3566) and cells were imaged with a Widefield Leica microscope DMI6000B.

### *In-vivo* photo-ablation using injection or micro emulsion as delivery system

The skin at the nape of the neck of wildtype and IL31RA^-/-^ mice was shaved and injected with 5 μM IL31^SNAP^ or IL31^K138A-SNAP^ coupled to 15 μM IR700 in a 50 μl volume. 20 minutes after the injection, near infra-red light (680nm) at 200 J/cm^2^ was applied on the injection site for 2 minutes. This procedure was repeated for 3 consecutive days. For von Frey and Hot Plate tests, this procedure was performed on the hind paw using a 20 μl injection volume. All the behavioral experiments were performed 1 week after the last injection.

The micro emulsion was prepared as already described ^36^ Briefly, all the components were assembled as follow: Caprylic Triglyceride 81gr; Glyceryl Monocaprylate 27gr; Polysorbate80 12gr; Sorbitan Monooleate 8gr. The micro emulsion was mixed with the coupling reaction (IL31^K138A-SNAP^ + IR700) at 1:1 ratio in 10 μl volume with 5 μM as IL31^K138A-SNAP^ final concentration. The mixture was then applied on Calcipotriol-treated mice and FPLCA mice following the same procedure described above for the injection.

### Immunofluorescence

Back skin from 8-10 week-old Black 6/J male mice was dissected and fixed in PFA 4% overnight at 4°C. The next day, skin samples were transferred in 30% sucrose for 3-4 hours and frozen in OCT. 40-μm sections were cut with a cryostat and stained for Rabbit anti-Keratin14 (K14) (Covance # PRB155P) or Rabbit anti-Pgp9.5 (Dako # Z5116) antibodies (1:200 dilution) in PBS containing 5% goat serum + 0,3% Triton-X overnight at 4°C. Secondary anti-Rabbit Alexa488 antibody was diluted 1:1000 and incubated for 2 hours at 4°C in the dark. 1 μg/ml Dapi (Invitrogen # D1306) was used to stain nuclei. Slides were mounted with Prolong gold antifade (Invitrogen # P36930) and imaged with a Leica SP5 confocal microscope and analyzed with ImageJ. To quantify K14+ cell, the number of K14+ per Dapi^+^ cells in the epidermal basal layer was considered and expressed as %. For Pgp9.5+ fibers quantitation, the region of dermis / epidermis junction was analyzed and only the intra-epidermal free nerve endings per section were considered for counting ^66^

### Immunohistochemistry

Paraffin-embedded back skin from 8-10 week-old Black 6/J or FPLCA male mice was sectioned at 6μm thickness with a microtome. Sections were deparaffinized, hydrated to distilled water and stained with Hematoxylin and Eosin for morphology and cellular infiltration, Toluidine blue for mast cells and Congo Red (Bio-optica # 04210822) for amyloid deposits. Slides were mounted and imaged with a Microdissector Leica microscope LMD7000.

### Flow cytometry

To analyze differences in Langerhans cell levels upon photoablation, we performed flow cytometry. The whole back skin of wildtype 8-10 week-old Black 6/J male mice was collected at different time points post photoablation (1 day, 1 week, 1 month post) and cutaneous dendritic cells were isolated as previously described ^67^ Briefly, the back skin was cut in 1cm pieces and digested overnight at 4° C in 1 mg/ml Dispase II (Roche # 4492078001). The next day, the skin pieces were further digested in an enzymatic solution of 1 mg/ml Collagenase IV (Worthington # CLS4 LS004188) and 0,5 mg/ml DNase (Sigma # DN25) at 37°C for 2 hours; finally, the cell suspension was filtered with a 100 μm cell strainer, washed with PBS + 0,5 M EDTA + 2% FBS and centrifuged at 1700 rpm for 5 minutes at 8°C. Cells were re-suspended in HBSS + 5mM MgCl2 + 100 μg/ml DNase + 1% BSA and labelled with 0, 5 μg /1x10^6^ cells PE anti-mouse CD207 (Langerin) (Biolegend # 144204) or 0,25 μg /1x10^6^ cells Alexa Fluor 488 anti mouse IA/IE (MHCII) (Biolegend # 107616) antibodies, or Dapi (1 μg/ml) in PBS + 2% BSA. Langerhans cells were distinguished by the other cellular populations for their double expression of Langerin and MHCII ^67, 68^ The experiments were performed using FACS Aria instrument and data were analyzed using FlowJo software.

### SDS-Page and Western blot

To assess the coupling reaction, 1 and 2 μM IL31^K138A-SNAP^ was coupled with 3 and 6 μM BG549 respectively for 1 hour at 37°C. The coupling reactions were analyzed by SDS-PAGE on a precast acrylamide gel (BioRad #456-9034), along with the same concentrations of IL31^K138A-SNAP^ alone. The bands corresponding to the binding of IL31^K138A-SNAP^ with BG549 were visualized by gel fluorescence. All the samples were visualized by Coomassie staining. For IL31-mediated signaling, back skin from wild-type mice were injected with vehicle (PBS), 5μM IL31^SNAP^ or IL31^K138A-SNAP^. After 1 hour, mice were sacrificed and skin was collected and lysed in Ripa Buffer (Sigma, #R0278) with proteases inhibitor cocktail (Roche #11873580001). Protein lysates were quantified by Bradford assay. 30 μg total lysate were separated on 10% SDS-Page gel and transferred to a nitrocellulose membranes (Protran #10600007). Membrane were incubated with the following antibodies: anti STAT3 (Cell Signaling #9139), anti phospho STAT3 (Tyr705) (Cell Signaling #9131), anti MAPK (Cell Signaling #4695), anti phospho MAPK (Thr202/Tyr204) (Cell Signaling #9106), anti AKT (Cell Signaling #4691), anti phospho AKT (Ser473) (Cell Signaling #9271), anti-Actin (Cell signaling #4970). Bands were visualized using the ECL detection system (Amersham #RPN2106); band density was calculated using ImageJ and the levels of phosphorylated proteins were normalized to the total counterpart.

### Behavioral testing

#### Scratching behavior

To evaluate the scratching response, 8-10 week-old male wild-type or IL31RA^-/-^ or IL31RA^S476F^ mice were shaved at the nape of the neck, placed in Plexiglas chambers to acclimatize (30 minutes), videotaped for 30 minutes and scratching bouts were counted. One bout was defined as an event of scratching lasting from when the animal lifted the hind paw to scratch until it returned it to the floor or started licking it. For IL31-mediated itch, 5μM IL31^SNAP^ or IL31^K138A-SNAP^ or native IL31 (Peprotech, #210-31) were injected intradermal for 1-3 days depending on the experiment. Regarding other itch-mediators, 10 mM histamine (Sigma, #H7250), 1 mM LY344864 (Abcam, #ab120592) or 12,5 mM Chloroquine (Sigma, #C6628) were injected into the back skin of wild type mice previously injected with 5 μM IL31^K1^ 38^a-snap^ + 15 μM |R700 with or without near IR illumination. The injection of pruritogens was done 1 week after the last treatment. For long-term reversal IL31 evoked scratching, IL31 was injected 1 day after the last day of illumination and then every week for 8 weeks. 4μg/ml Calcipotriol-mediated itch was assessed at different time points to monitor dermatitis development, with or without near IR illumination.

#### von Frey test

Mice were habituated on an elevated platform with a mesh floor for 30 minutes. The plantar side of the hind paw was stimulated with calibrated von-Frey filaments (North coast medical #NC12775-99) to assess baseline levels of mechanical sensitivity. The hind paw of wildtype mice were then injected for 3 consecutive days with IL31 ^K1^38^A-SNAp^ coupled with IR700 with or without near IR light illumination. The stimulation with von-Frey filaments was then repeated 1 week after the last photoablation. The 50% paw withdrawal thresholds were calculated using the Up-Down method ^69^

#### Hot Plate test

Mice were injected for 3 consecutive days with IL31^K138A-SNAP^ coupled with IR700 with or without near IR light illumination. 1 week after the last injection, mice were placed on top of a Hot Plate (Ugo Basile #35150) that was preset to 52¼C and the latency to response as distinguished by flicking or licking of the hind paw was observed. In order to avoid injury to the mice, a cutoff of 30 seconds was set.

### Mouse model for Atopic Dermatitis (AD)

4 μg/ml of the vitamin D3 analogue Calcipotriol (Tocris #2700) was applied topically to the shaved back skin of the mice for 14 or 21 days, depending on the experiment. 0,002% DMSO was applied as vehicle control for 14 days. Atopic dermatitis development was assessed by scratching behavior over a 30 minute-recording period, skin thickness measurement using a Caliper and histological analysis for cellular infiltration of the treated skin by Hematoxylin & Eosin staining.

### Mouse model for Familiar Primary Localized Cutaneous Amyloidosis (FPLCA)

The IL31RA^S476F^ mouse was generated using cloning-free CRISPR/Cas9 technology as previously described ^70^ Briefly, the knock-in point mutation in Il31ra locus, responsible for the amino acid substitution S476F ^26^, was obtained by zygote pronuclear injection of a mixture of 30 ng/μl Cas9 protein (Produced and purified by EMBL Protein Expression and Purification Core Facility), 0,61 pmol/ μl IL31RA-crRNA (IDT technology), 0,61 pmol/ μl tracrRNA (IDT technology) and 10 ng/μl single strand oligo DNA (ssODN) donor carrying the desired point mutation (C>T) (IDT technology). A 120 nucleotides long ssODN was designed containing two homology arms of 60 nucleotides each, flanking the mutation site. Additional silent point mutations were included in the donor sequence to avoid Cas9-mediated cut. Sequences are listed in supplementary figure 4.

### Mouse model for psoriasis

62,5 mg Aldara 5% corresponding to 3,125 mg of active component was applied topically to the back skin for 7-10 days as previously described ^71^. The development of the disease was monitored and graded using a modified human scoring system Psoriasis Area Severity Index (PASI) which considers skin thickness, erythema and scaling. Scratching behavior was also recorded. After 7-10 days, mice were sacrificed and skin was collected for histological analysis.

### Nerve photoablation

The sciatic nerve innervating the hind paw/calf region of 8-10 week-old male mice was treated with 30 ng/μl TPA to permeabilize the epineural sheath for 15 minutes; then 5 μM IL31 ^K138a-snap^+1 5 μM |R700 was applied on the exposed nerve and after 10-15 minutes, near IR light was applied for 45 seconds, except in the control group. Starting from 3 days post treatment, mice were injected with 5 μM IL31^SNAP^ at different days in the paw/calf level, and monitored for scratching expressed as time spent in biting at the injected area.

### Statistical analysis

All statistical data are presented as Standard error of the mean (SEM) along with the number of samples analyzed (n). Student’s t-test and/or analysis of variance ANOVA followed by the appropriate postHoc test were used. Statistical significance was assumed at p<0.05.

